# Tasman-PCR: A genetic diagnostic assay for Tasmanian devil facial tumour diseases

**DOI:** 10.1101/287847

**Authors:** Young Mi Kwon, Maximilian R. Stammnitz, Jinhong Wang, Kate Swift, Graeme W. Knowles, Ruth J. Pye, Alexandre Kreiss, Sarah Peck, Samantha Fox, David Pemberton, Menna E. Jones, Rodrigo Hamede, Elizabeth P. Murchison

## Abstract

Tasmanian devils have spawned two transmissible cancer clones, known as devil facial tumour 1 (DFT1) and devil facial tumour 2 (DFT2). DFT1 and DFT2 are transmitted between animals by the transfer of allogeneic contagious cancer cells by biting, and both cause facial tumours. DFT1 and DFT2 tumours are grossly indistinguishable, but can be differentiated using histopathology, cytogenetics or genotyping of polymorphic markers. However, standard diagnostic methods require specialist skills and equipment and entail long processing times. Here, we describe Tasman-PCR: a simple PCR-based diagnostic assay that identifies and distinguishes DFT1 and DFT2 by amplification of DNA spanning tumour-specific interchromosomal translocations. We demonstrate the high sensitivity and specificity of this assay by testing DNA from 544 tumours and 818 normal devils. A temporal-spatial screen confirmed the reported geographic ranges of DFT1 and DFT2 and did not provide evidence of additional DFT clones. DFT2 affects disproportionately more males than females, and devils can be co-infected with DFT1 and DFT2. Overall, we present a PCR-based assay that delivers rapid, accurate and high-throughput diagnosis of DFT1 and DFT2. This tool provides an additional resource for devil disease management and may assist with ongoing conservation efforts.

## Introduction

Tasmanian devils *(Sarcophilus harrisii)* are marsupial carnivores endemic to the Australian island of Tasmania. This species is affected by two transmissible cancers, known as Tasmanian devil facial tumour 1 (DFT1) and Tasmanian devil facial tumour 2 (DFT2). DFT1 and DFT2 manifest as facial tumours collectively known as devil facial tumour disease (DFTD) and are transmitted between animals by the direct transfer of living allogeneic cancer cells, probably by biting (Pearse and Swift, 2006; Pye et al., 2016). Thus, DFT1 and DFT2 are two independent malignant clones, one of which arose from the somatic cells of an individual female devil (DFT1), the other from a male (DFT2), and which both now propagate through the devil population via transfer of contagious cancer cells (Deakin et al., 2012; Murchison et al., 2012; Pye et al., 2016). DFT1 was initially reported in 1996 in north-east Tasmania and is now widespread across the island, causing significant declines in devil populations (Hawkins et al., 2006; Lazenby et al., 2018; McCallum, 2008). DFT2, on the other hand, which was first observed in 2014, has been reported in only five animals, all located in the Channel, a ~550 km^2^ peninsula in Tasmania’s south-east (Pye et al., 2016).

DFTD tumours caused by DFT1 and DFT2 are grossly indistinguishable, but are histologically, cytogenetically and genetically distinct (Pye et al., 2016; Stammnitz et al., 2018). Differential diagnosis of DFT1 and DFT2 is routinely performed with histopathology and immunohistochemistry (Loh et al., 2006; Murchison et al., 2010; Pye et al., 2016; Tovar et al., 2011). Diagnosis can also be confirmed using cytogenetics, as DFT1 and DFT2 have markedly different karyotypes (Pearse and Swift, 2006; Pye et al., 2016). Genetic methods, based on microsatellite and major histocompatibility complex (MHC) genotypes, have also been used to distinguish DFT1 and DFT2 (Pye et al., 2016). However, the presence of host tissue within DFT1 and DFT2 tumour biopsies, as well as allelic variation across tumour and host populations, often leads to ambiguity with interpretation of such genotypes.

DFT1 and DFT2 are foreign tissue grafts in their hosts and must escape the allogeneic immune system. DFT1 cells down-regulate MHC class I, perhaps contributing to this clone’s low immunogenicity (Siddle et al., 2013); the mechanism of DFT2 immune avoidance is unknown. Interestingly, however, whereas DFT1 equally affects males and females (Loh et al., 2006), all five devils diagnosed with DFT2 have been male (Pye et al., 2016).

DFT1 was first detected in the Channel in 2012, shortly before the discovery there of DFT2 in 2014 (Pye et al., 2016). Thus, DFT1 and DFT2 have overlapping host ranges within the Channel, and in the period from December 2012 to June 2015, seven cases of DFT1 and five of DFT2 were reported in the area (Pye et al., 2016). While it is known that a devil can carry two strains, or subclones, of DFT1, derived from different infections (Murchison et al., 2012), it remains unknown whether a host can possess both DFT1 and DFT2 tumours simultaneously.

Transmissible cancers have been rarely observed in nature. There are currently eight known naturally occurring transmissible cancers, five of which affect various species of bivalve molluscs (Metzger et al., 2015; Metzger et al., 2016; Murchison, 2008; Pye et al., 2016). In 1996, when DFT1 was first observed, only one other example was known, a sexually transmitted venereal tumour in dogs (Murchison, 2008). Furthermore, there was no evidence of devil tumours similar to DFTD prior to 1996 (Hawkins et al., 2006; Loh et al., 2006). Thus, the discovery of DFT2 was surprising, and suggests that Tasmanian devils may be particularly vulnerable to the emergence of transmissible cancers (Stammnitz et al., 2018). There is a possibility that additional DFT clones may or may have existed in the devil population, but escaped detection. Furthermore, the extent of the DFT2 range beyond the Channel is not yet confirmed.

The identification of two transmissible cancers within an interval of eighteen years raises important questions about Tasmanian devils and their susceptibility to transmissible cancers. Here, we describe the design and validation of a genetic diagnostic test for confirmation of DFT1 and DFT2 in devils. We use this “Tasman-PCR” test to examine the range of DFT2 and to confirm the absence of additional DFT clones within a cohort of 544 tumours collected between 2003 and 2016 from more than 69 locations around Tasmania. Of the eleven devils detected with DFT2 (including the five previously reported cases), nine are male, suggesting that females may have reduced susceptibility to this clone. Furthermore, we identified two individual devils simultaneously hosting DFT1 and DFT2 tumours. Importantly, this work provides a diagnostic tool for rapid and high-throughput assessment of DFT status and may assist with devil disease monitoring and conservation efforts.

## Results

### Development of PCR-based markers for DFT1 and DFT2

DFT1 and DFT2 are both clonal lineages, hence somatic variants that were acquired early during each lineage’s history may present useful markers for PCR-based diagnostic assays. Interchromosomal translocations are particularly effective markers, as the presence or absence of an amplification product is the screening end-point. We assessed a set of interchromosomal structural variants that had been identified and validated in two DFT1 and in two DFT2 genomes respectively, but not found in 34 normal devil genomes (Stammnitz et al., 2018), and selected two putative DFT 1-specific markers and two putative DFT2-specific markers for further screening (Table 1). A triplex PCR assay was then optimised, containing primers specific for a DFT1 marker (DFT1-A, 231 base pairs, bp), a DFT2 marker (DFT2-A, 321 bp), as well as positive control primers amplifying the endogenous *RPL13A* locus (520 bp) (Siddle et al., 2013) (Figure 1).

**Figure 1:**
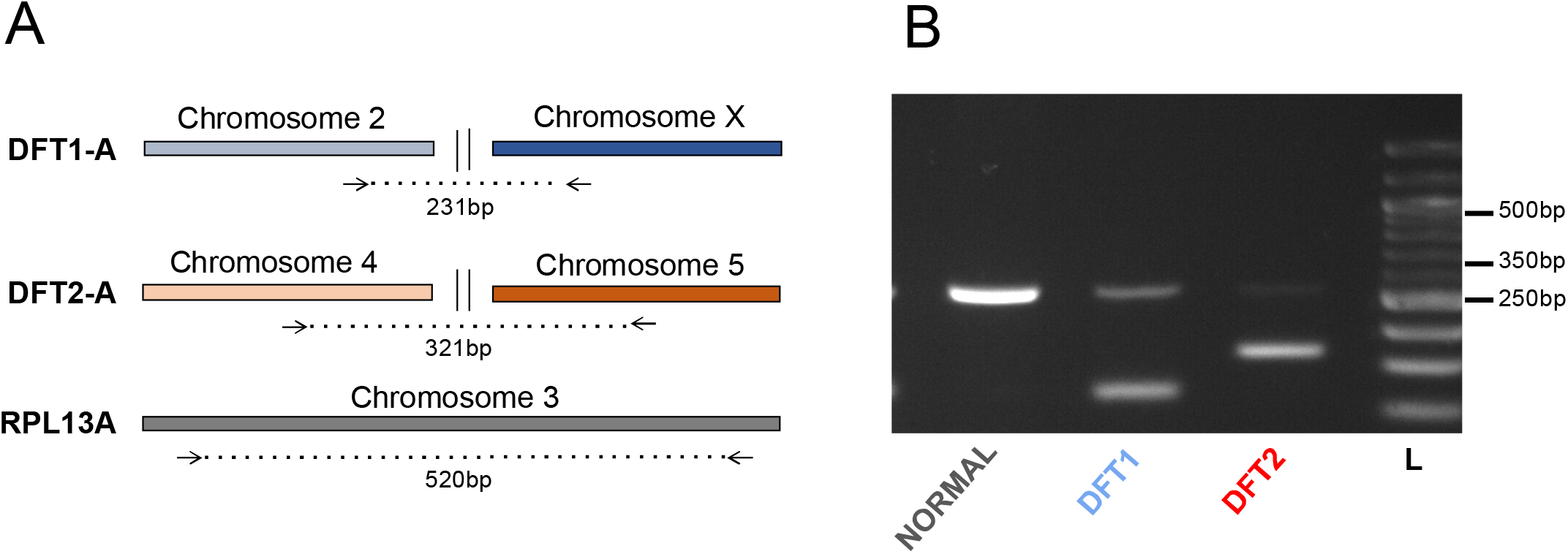
Tasman-PCR, a DFTD triplex PCR. (**A**) Design of triplex DFTD PCR using primers spanning DFT1-A, DFT2-A and *RPL13A.* (**B**) Gel electrophoresis showing amplification products from normal devil DNA, as well as from DFT1 and DFT2 DNA. L, ladder.

**Table 1:**
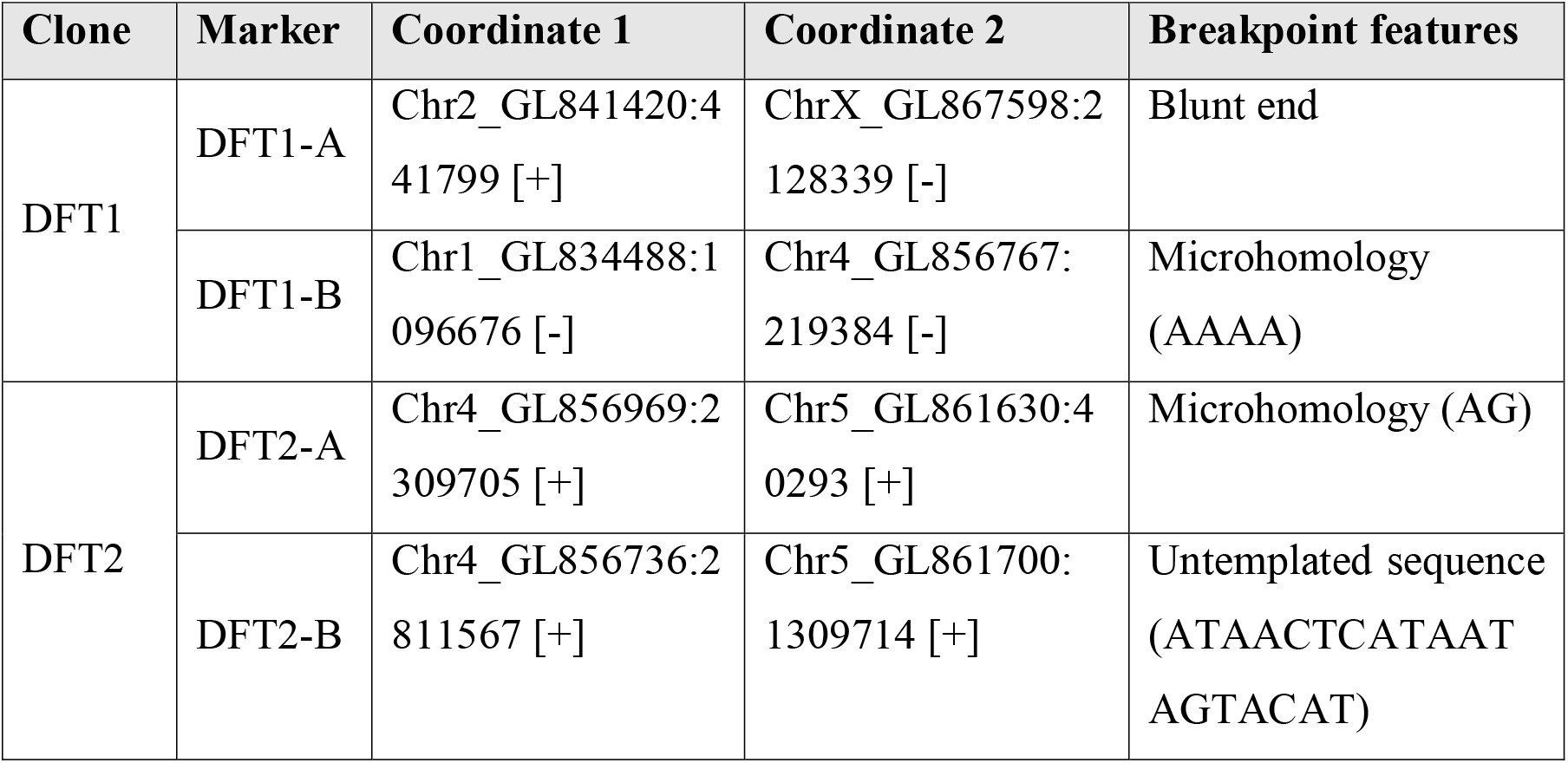
Coordinates and breakpoint features for structural variants specific to DFT1 and DFT2 used in screen. Coordinates are relative to the Tasmanian devil reference genome Devil_refv7.0 (Murchison et al., 2012).

### Normal devil screen

Interchromosomal translocations are more common in cancer than in germline DNA. However, it is possible that DFT1-A and/or DFT2-A were in fact inherited polymorphisms present in the constitutive genomes of the individual devils that spawned the respective clones, and hence may also be found in the genomes of some normal devils in the population. In order to assess this, we screened our triplex PCR assay across normal DNA from 818 devils, including both nondiseased individuals as well as devils infected with DFT1 and/or DFT2, sampled between 2003 and 2016 from more than 64 locations in Tasmania (Table S1). DNA from three devils (0.4%) was faintly positive for DFT1-A, and no devils amplified DFT2-A (Table S1). We further assessed the three DFT1-A-positive DNA samples by amplifying a panel of microsatellite length polymorphisms. All three samples amplified three or more alleles at one or more loci, suggesting DNA contamination (Table S2A). Furthermore, all three DNA samples which were positive for DFT1-A also amplified DFT1-B (Table 1), further suggesting the possibility that these normal samples may have been contaminated by DFT1 DNA. Altogether, these results suggest that the tumour DNA markers used in our triplex assay are highly specific for DFT1 and DFT2, and are likely to be somatic translocations that arose subsequent to neoplastic transformation in these two clones.

### Analysis of confirmed DFTD tumours

We next used the triplex assay to screen 342 confirmed DFTD (DFT1 and DFT2) tumours (Table S3A). The tumours were collected between 2003 and 2016 from 56 locations in Tasmania, and DFTD status was confirmed through histopathology, cytogenetics and/or microsatellite genotyping (Murchison 2010, Loh 2006, Pearse 2012). 329 of 342 (96.2%) confirmed DFTD tumours were positive for either DFT1-A or DFT2-A (321 DFT1-A positive, 8 DFT2-A positive), and no tumours were positive for both DFT1-A and DFT2-A (Table S3A).

We considered the following four explanations for the 13 confirmed DFTD tumours which did not amplify DFT1-A or DFT2-A (Table S3A): (1) they belong to a DFT1 or DFT2 subclone which diverged from a clonal ancestor prior to DFT1-A or DFT2-A being acquired; (2) they belong to a DFT1 or DFT2 subclone which lost DFT1-A or DFT2-A, or acquired mutations within the DFT1-A or DFT2-A primer binding sites; or (3) the biopsy used for DNA extraction included only non-neoplastic host tissue, or included DFTD cells at levels that were undetectable under our PCR and gel electrophoresis conditions. To distinguish between these three possibilities, we first screened the 13 tumours with DFT1-B and DFT2-B (Table 1); none of these samples amplified either marker. Next, we screened the 13 samples, together with their matched hosts, across a panel of four polymorphic microsatellites (Table S2B). None of the 13 samples showed the characteristic microsatellite profiles of DFT1 or DFT2 (Pye et al., 2016), suggesting that explanations (1) and (2) above are unlikely to apply. Furthermore, all 12 of these samples for which matched host tissue was available showed microsatellite profiles which were identical to those of their hosts (Table S2B). Thus, it appears likely that the biopsies used for DNA extraction carried only host DNA, or that DFTD DNA was present at a low level and was undetectable under our assay conditions (explanation (3) above). To further assess this possibility, we performed an additional DNA extraction on one of these tumours and confirmed DFT1-A positivity on this second attempt. Thus, variation in contribution of neoplastic cells to tumour tissues should be considered in the event of an unexpected negative finding.

### Tumour screen

We used Tasman-PCR to screen DNA from 175 suspected DFTDs collected from 114 devils. In these cases, DFTD was suspected due to gross morphology and clinical presentation, but no records of confirmed diagnosis based on histopathology, cytogenetics or microsatellite genotyping were found. 158 of these suspected DFTDs were positive for DFT1-A (90.3%) and five were positive for DFT2-A (2.9%) (Table S3). None of the 12 DFT1-A and DFT2-A negative tumours were positive for DFT1-B or DFT2-B. Microsatellite genotyping confirmed that all nine DFT1-A and DFT2-A negative suspected DFTDs for which microsatellites could be amplified and for which matched host tissue was available were identical to their matched host (Table S2C). This suggests that these samples may be DFTD tumours from which only non-neoplastic host tissue was sampled for DNA extraction (explanation (3) above); alternatively, it is possible that these samples may be non-DFTD host-derived lesions.

We next screened DNA from 27 confirmed or suspected non-DFTD lesions. These lesions included cutaneous lymphomas, papillomas, squamous cell carcinomas and other carcinomas (Table S3C). Microsatellite alleles were analysed in the 27 non-DFTD samples, together with their matched hosts when available. Several samples had one or two microsatellite alleles which were not detectable within the tissues of the matched host (Table S2D). However, non-DFTD lesions from different individuals had different genotypes. This suggests that the tumour-unique microsatellite alleles arose via somatic mutation of matched host alleles and does not support the possibility that any of these 27 samples represent additional transmissible cancers.

### Distribution of DFT1 and DFT2

Of the 492 confirmed or suspected DFTD tumours which amplified a DFT1 or DFT2 tumour marker in Tasman-PCR, 13 tumours from 11 devils were positive for DFT2-A (Table S3). These DFT2-A-positive animals were all sampled in the Channel Peninsula between 2014 and 2016 (Figure 2). The remaining 479 tumours, collected from 63 locations between 2003 and 2016, amplified DFT1-A (Figure 2). Although we cannot rule out the possibilities that the current or historical DFT2 range extends beyond the Channel Peninsula, or that DFT2 arose prior to 2014, our results support the idea that DFT2 may have arisen recently within the Channel Peninsula.

**Figure 2:**
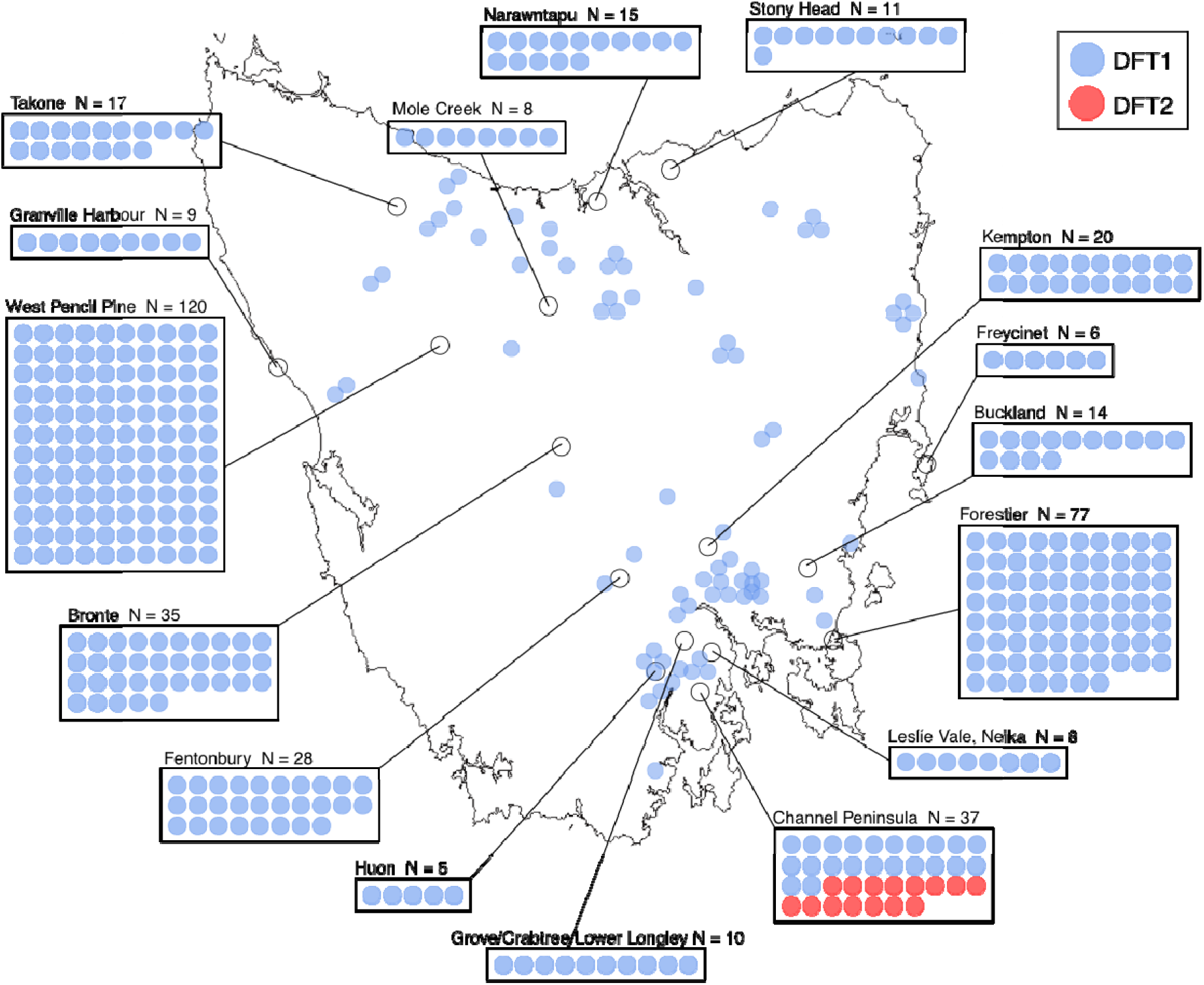
Distribution of DFT1 and DFT2. Locations of 492 tumours, sampled from 358 devils between 2003 and 2016, which amplified either DFT1-A (479 tumours) or DFT2-A (13 tumours). Each tumour is represented by a blue (DFT1) or red (DFT2) dot. The Channel Peninsula, where all DFT2-A-positive tumours were found, is located in Tasmania’s southeast.

### DFT1 and DFT2 host gender

Our screen detected 11 animals with DFT2 tumours, including the five previously reported cases (Pye et al., 2016). Of these, 9 (82%), were male (Fisher’s exact test for no difference in DFT2 host gender, p = 0.033) (Figure 3). In contrast, DFT1 tumours were equally distributed between male and female hosts (of the 345 animals with one or more DFT1 tumours with known gender, 165 (47.8%) were male and 180 (52.2%) were female, (Fisher’s exact test for no difference in DFT1 host gender, p = 0.451), as previously reported (Loh et al., 2006) (Figure 3). This skew towards male hosts in DFT2 suggests that this clone may preferentially affect males, although larger numbers of DFT2 cases will be required to confirm this.

**Figure 3:**
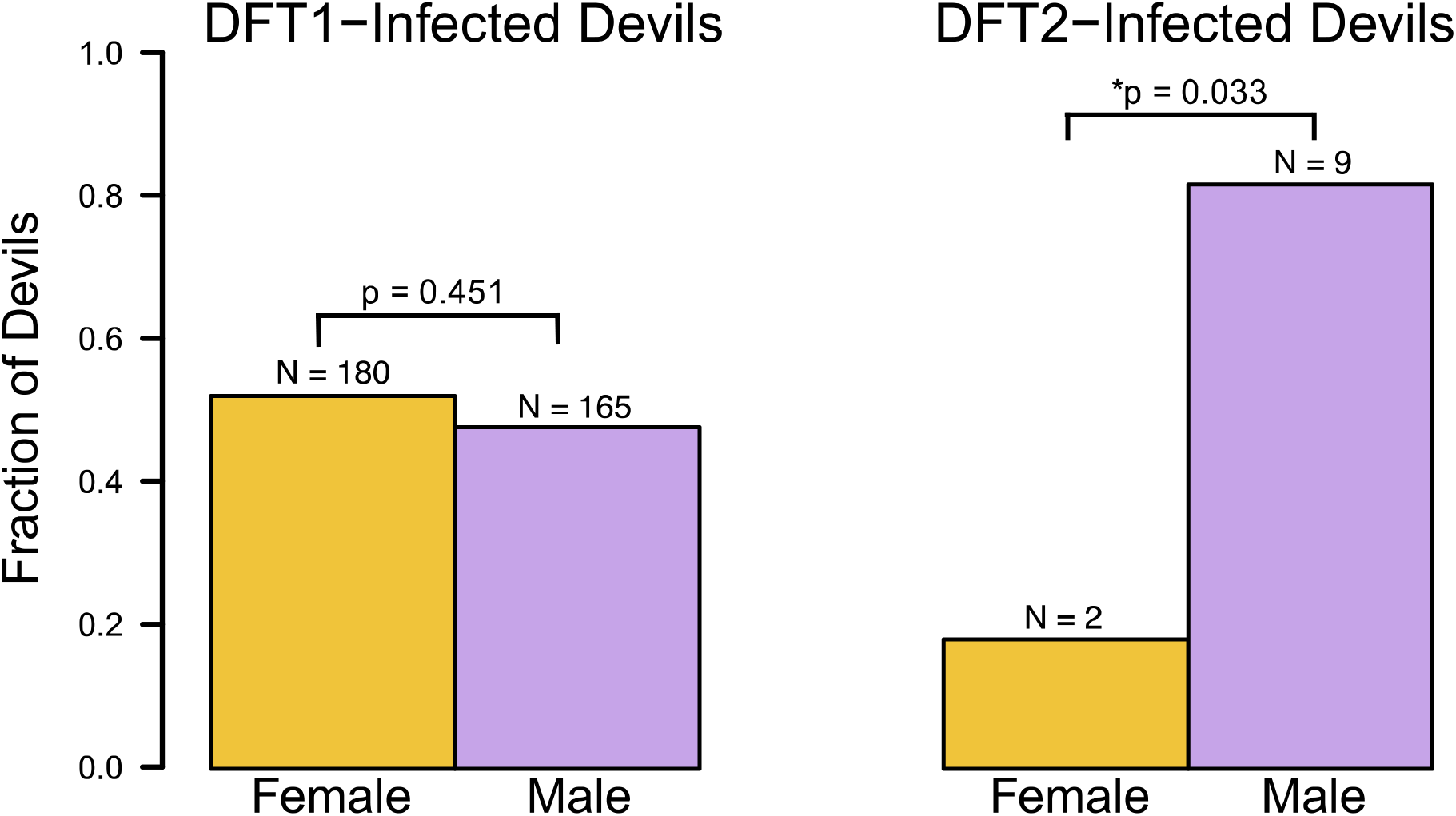
Gender distribution of DFT1 and DFT2. Gender distributions of 345 devils with one or more DFT1 tumours and 11 devils with one or more DFT2 tumours. ‘Fraction of devils’ denotes the fraction of devils hosting at least one DFT1 tumour that are male and female (left) or the fraction of devils hosting at least one DFT2 tumour that are male and female (right). p values represent outcome of Fisher’s exact test for no difference in DFT1 and DFT2 host gender and * denotes significance (p < 0.05).

### Co-infection with DFT1 and DFT2

It has previously been noted that individual devils can simultaneously harbour tumours belonging to two distinct strains, or subclones, of DFT1 (Murchison et al., 2012). As the DFT1 and DFT2 ranges overlap in the Channel Peninsula, we investigated the possibility of coinfection with DFT1 and DFT2. We identified two individuals, Devil 812 and Devil 818, which carried both DFT1 and DFT2 tumours (Figure 4). Devil 812 was a four-year old female with three facial tumours, denoted 812T1, 812T2 and 812T3. 812T2 and 812T3 were both DFT1, whereas 812T1 was DFT2 (Figure 4). Devil 818, a two-year old male, carried a large DFT2 tumour, 818T1, ventral to the left ear, as well as a DFT1 tumour on the hard palate, 818T2 (Figure 4). Devil 818 also had a DFT2 mass in the left pre-auricular lymph node (818T3) (Figure 4), likely a metastasis from 818T1.

**Figure 4:**
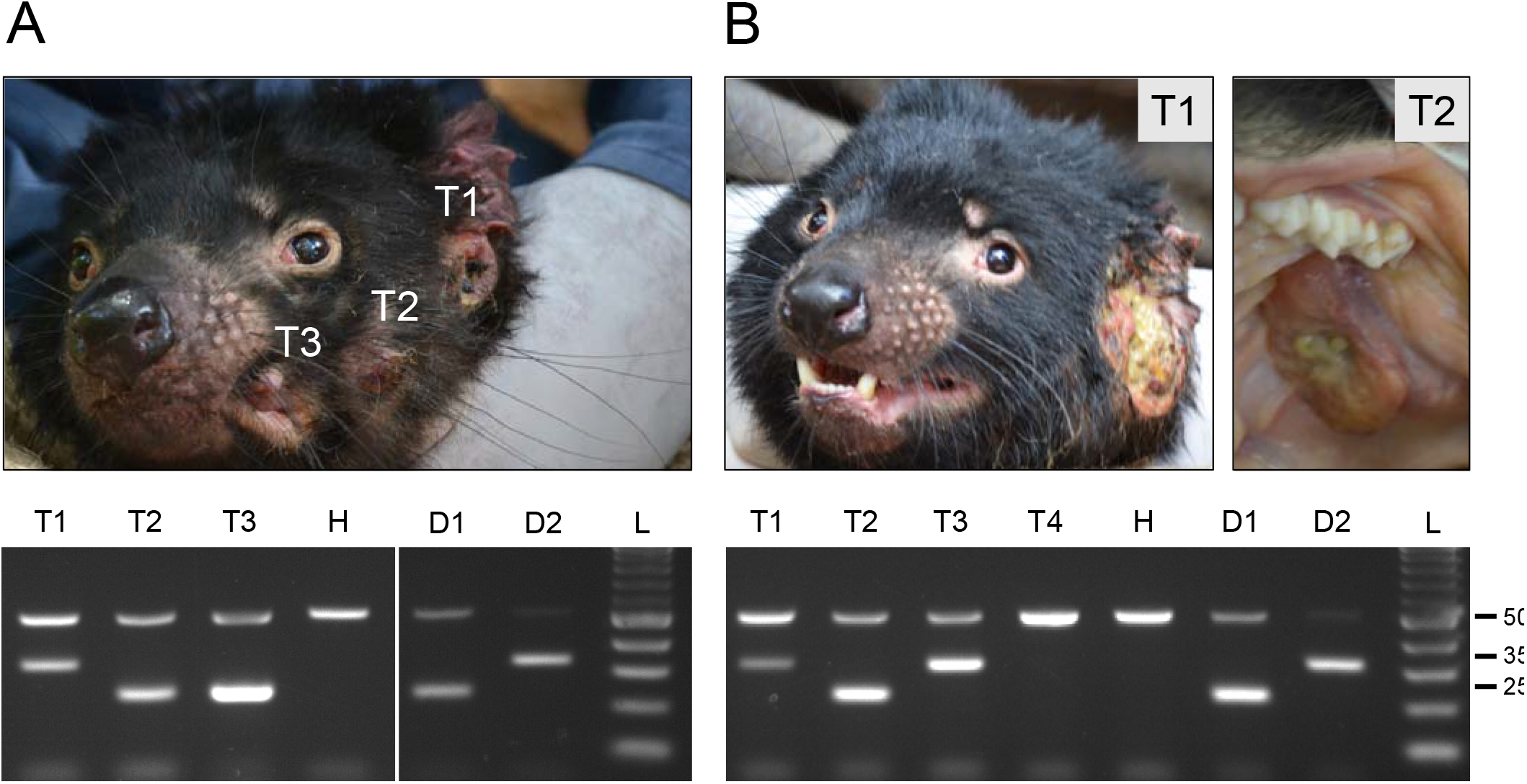
Co-infection with DFT1 and DFT2. Two devils were identified with both DFT1 and DFT2 tumours, **(A)** Devil 812 and **(B)** Devil 818. T1, T2, T3, T4 denote individual tumours from each animal, some of which are depicted in photographs. PCR results from each tumour are shown on gel and H denotes normal DNA from matched host. D1, D2 and L denote DFT1 control, DFT2 control and ladder respectively. In Devil 818, T3 was a tumour involving the left pre-auricular lymph node and T4 was a suspected tumour found in the left submandibular lymph node.

## Discussion

We present a PCR-based genetic diagnostic test for DFTD, Tasman-PCR, which complements existing diagnostic methods to provide rapid and cost-effective confirmation of DFT1 and DFT2. The test uses amplification of tumour-specific interchromosomal translocations to identify DFT1 and DFT2, and to distinguish between DFTD (DFT1 or DFT2) and non-DFTD tumours in devils. We used the assay to screen 544 devil tumours collected from more than 69 locations in Tasmania between 2003 and 2016.

Although the assay that we describe provides a useful method for diagnosis of DFT1 and DFT2, it has some limitations. Users should be aware of the potential for DNA cross-contamination to produce false positive results, and electrophoresis-based detection methods have limited sensitivity in samples with very low tumour cell fractions. DFT1 and DFT2 tumours are heterogeneous tissues which contain both neoplastic and normal cell components. While very little input material is required for PCR, which allows for minimally invasive, small biopsies, it can also increase the likelihood of obtaining biopsies that contain no neoplastic cells, leading to false negative results. Furthermore, although we did not find evidence of germline DFT1-A or DFT2-A in a screen of 818 normal devils, we cannot exclude the possibility that these markers are rare polymorphisms in devil populations; we suggest that users analyse matched normal tissues alongside tumours to control for this possibility. Additionally, although our screen did not identify any clear cases of DFT1 or DFT2 tumours negative for DFT1-A or DFT2-A respectively, it is possible that such lineages may have previously arisen or may occur in future. Users encountering unexpected negative results should perform PCR for additional markers, such as DFT1-B and DFT2-B (Table 1) as well as repeating DNA extraction and PCR from a separate biopsy of the same tumour. Importantly, this assay is designed to be complementary with other established DFT1 and DFT2 diagnostic methods, such as histology and cytogenetics, while being rapid, scalable, and minimally invasive. Further, it would be possible to develop this method to obtain real-time diagnosis in the field with the use of portable PCR laboratories (Koo et al., 2013; Taylor et al., 2014).

The discovery of two transmissible cancers in Tasmanian devils raises the possibility that additional, so far undetected, transmissible cancer clones exist within devil populations. In this study, we screened 544 Tasmanian devil tumours, collected between 2003 and 2016 from more than 69 locations in Tasmania (Table 2). 479 (88.1%) were DFT1 and 13 (2.4%) were DFT2; the remaining 52 (9.6%) were either confirmed or suspected host-derived non-DFTD lesions (27, 5.0%) or biopsies that shared identical microsatellite profiles with their matched hosts (as well as four biopsies for which matched host tissue was unavailable or microsatellites could not be amplified) (25, 4.6%) (Table 2). Thus, this screen has not detected evidence for additional transmissible cancers in Tasmanian devils. However, the method provides a useful tool for devil tumour monitoring and surveillance.

**Table 2:** Results of screen of 544 Tasmanian devil tumours. ‘No information on similarity to matched host’ denotes tumours for which matched host DNA was unavailable or for which microsatellite loci failed to amplify.

| Marker amplified | Number of tumours |
| --- | --- |
| DFT1-A | 479 |
| DFT2-A | 13 |
| *RPL13A* only | 52<br>27 Non-DFTD lesions<br>21 Microsatellite alleles identical to matched host<br>4 No information on similarity to matched host |
| Total | 544 |

DFT1 had been detected in the Channel Peninsula prior to the discovery there of DFT2, and previous studies have shown that different DFT1 strains, or subclones, can co-infect one host (Murchison et al., 2012). Here, we confirmed that DFT1 and DFT2 can also co-infect the same devil. Co-infections of these two phenotypically similar DFTs highlight the importance of careful monitoring, as competition for a limited host devil population may influence evolution of virulence (Alizon et al., 2009).

Of the 11 DFT2 cases that we detected, nine involved a male host (Fisher’s exact test for no difference in DFT2 host gender, p=0.033). This apparent preference for male hosts may reflect underlying transmission dynamics. Alternatively, it is possible that females are less susceptible to DFT2, perhaps due to recognition of Y chromosome-derived allogeneic antigens (Stammnitz et al., 2018). Nevertheless, we detected two females with DFT2 tumours, confirming that this disease does not exclusively affect males. The gender bias observed in DFT2 may impact devil population structures by reducing overall male devil abundance; over time, this may decrease reproductive success and increase inbreeding by limiting mating opportunities. Furthermore, if DFT2 is largely excluded from female hosts, this may affect the future potential for this clone to survive, perhaps selecting for the emergence of DFT2 subclones with balanced host gender proclivity through Y chromosome loss (Stammnitz et al., 2018).

Overall, this study presents Tasman-PCR, a PCR-based assay for rapid, high-throughput diagnosis of DFT1 and DFT2 in Tasmanian devils. By screening 544 tumours, we have documented the distributions of DFT1 and DFT2, and identify two devils co-infected with both clones. Our analysis does not provide evidence for additional DFTD clones beyond DFT1 and DFT2. The co-occurrence of two transmissible cancer clones in Tasmanian devils presents unique challenges for the conservation of this iconic species. The genetic diagnostic screening tools described here provide important additional resources for population monitoring and disease surveillance.

## Materials and Methods

### Animal ethics, sample collection and DNA extraction

Biopsies were collected from wild or captive Tasmanian devils or from animals whose carcasses were found as roadkill. Tumour and host (ear, liver, spleen, blood, gonad, or submandibular lymph node) tissue biopsies were collected into either RNAlater or ethanol. Fine needle aspirates collected into a 1:1 solution of Buffer AL (Qiagen) : PBS were sampled from tumors that were too small to biopsy. All animal procedures followed a Standard Operating Procedure that was approved by the General Manager of the Natural and Cultural Heritage Division in the Tasmanian Government Department of Primary Industries, Parks, Water and the Environment (DPIPWE), in agreement with the DPIPWE Animal Ethics Committee, or under University of Tasmania Animal Ethics Permits A0014976, A0010296, A0013326 and A0016789. The sample collection procedures were approved by the University of Cambridge, Department of Veterinary Medicine Ethics and Welfare Committee (CR191). A subset of tumours were previously confirmed as DFT1 or DFT2 using histopathology, cytogenetics or microsatellite genotyping (Pearse et al 2012, Loh et al 2006, Murchison 2010). Genomic DNA was extracted using the Qiagen DNeasy Blood and Tissue Kit. Genomic DNA was amplified using the Illustra GenomiPhi V2 DNA Amplification Kit (GE Healthcare) prior to PCR amplification.

### Tasman-PCR amplification and gel electrophoresis

Primers used in this study are as follows:

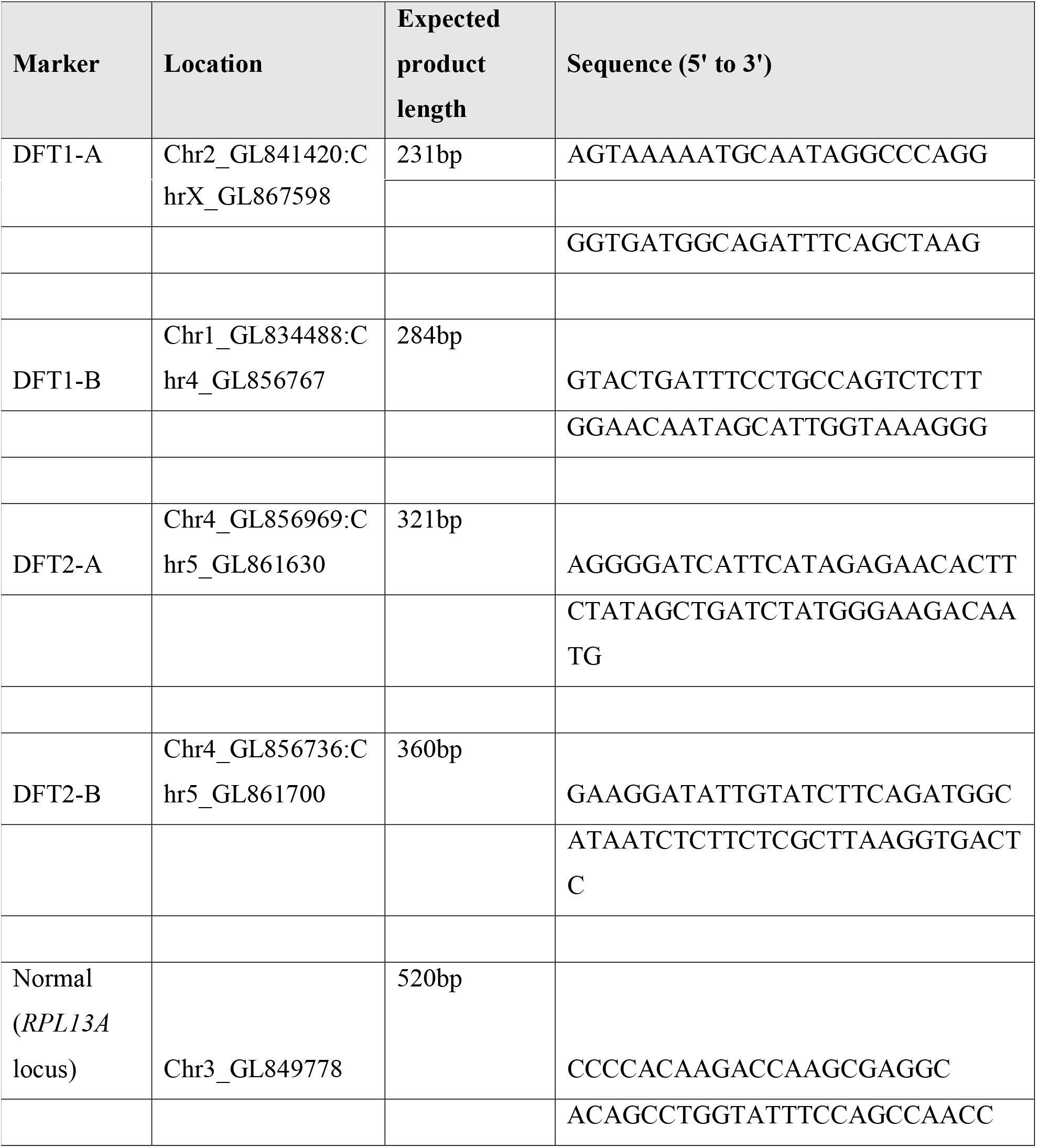

The triplex PCR was performed in 20 μl reactions with 2 units Qiagen Taq Polymerase, 1X CoralLoad PCR buffer, 0.2 mM each dNTP and ~20 ng genomic DNA per reaction. Primers were at final concentration of 0.4 μM (DFT1-A and DFT2-A primers) or 0.8 μM *(RPL13A* primers). Amplification was performed using a Nexus GS-1 Thermal Cycler (Eppendorf) with the following conditions:

| Temperature (°C) | Time | Cycles |
| --- | --- | --- |
| 94 | 3 min 30 s | 1 |
| 94 | 30 s | 30 |
| 60 | 30 s |  |
| 72 | 30 s |  |
| 72 | 5 min | 1 |

Single (non-triplex) PCRs, including those involving DFT1-B and DFT2-B, were performed with the same conditions, with primers at final concentration 0.4 μM. Gel electrophoresis was performed with a 3% agarose gel and products were detected using ethidium bromide.

### Microsatellite Genotyping

Microsatellite amplification and fragment size detection was performed as previously described (Pye et al., 2016) and analysed using the Fragman package (Covarrubias-Pazaran et al., 2016).

## Acknowledgements

This work was supported by grants from Wellcome (102942/Z/13/A), the National Science Foundation (DEB-1316549), the Australian Research Council (DE170101116), the University of Tasmania Foundation (Eric Guiler Tasmanian Devil Research Grants) and a Philip Leverhulme Prize awarded by the Leverhulme Trust. Y.M.K. is supported by a Herchel Smith Postgraduate Fellowship and M.R.S is supported by a scholarship from the Gates Cambridge Trust. We are grateful to the Save the Tasmanian Devil Program and to individuals who collected samples. In particular, we thank Nick Mooney, Billie Lazenby, Robyn Taylor, Anne-Maree Pearse and Tasha Czarny.

## Supplementary tables

**Table S1: Normal devil screen.**

DNA extracted from normal tissues from 818 devils was screened with the triplex PCR incorporating DFT1-A (D1), DFT2-A (D2) and *RPL13A* (N). Individual and sample ID, microchip (if available), location, gender and year-of-sampling are indicated for each individual, as well as information about markers amplified.

**Table S2: Microsatellite genotyping.**

(**A**) Microsatellite genotypes across four loci (C, D, F and N) from normal tissues from 3 devils which amplified DFT1-A. DFT1 and DFT2 genotypes are shown; two different DFT1 genotypes at locus C have been observed. Alleles which were weaker than expected, and are likely due to contamination, are marked with *.

(**B**) Microsatellite genotypes across four loci (C, D, F and N) from paired tumour and matched host tissues in 13 confirmed DFT1 tumours which failed to amplify DFT1-A and DFT1-B. DFT1 and DFT2 genotypes are shown; two different DFT1 genotypes at locus C have been observed. ‘-’ denotes matched host tissues that are unavailable.

(**C**) Microsatellite genotypes across four loci (C, D, F and N) from paired tumour and matched host tissues in 12 suspected DFT1 tumours which failed to amplify DFT1-A and DFT1-B. DFT1 and DFT2 genotypes are shown; two different DFT1 genotypes at locus C have been observed. ‘-’ denotes matched host tissues that are unavailable and ‘N/A’ denotes samples that failed to amplify microsatellite loci.

(**D**) Microsatellite genotypes across four loci (C, D, F and N) from paired tumour and matched host tissues in 27 confirmed or suspected non-DFTD tumours. DFT1 and DFT2 genotypes are shown; two different DFT1 genotypes at locus C have been observed. Alleles which were weaker than expected, which are likely to be due to mixed neoplastic and normal cell populations, are marked with *. ‘-’ denotes matched host tissues that are unavailable and ‘N/A’ denotes samples that failed to amplify microsatellite loci.

**Table S3: Tumour screen.**

DNA extracted from (A) confirmed DFTD tumours, (B) suspected DFTD tumours or (C) confirmed or suspected non-DFTD tumours was screened with Tasman-PCR incorporating DFT1-A (D1), DFT2-A (D2) and *RPL13A* (N). Individual and sample ID, microchip (if available), location, gender and year-of-sampling are indicated for each tumour, as well as information about markers amplified. Suspected or confirmed diagnosis is indicated for non-DFTD tumours. Different tumours collected from the same individual devil share the same individual ID but have unique sample IDs. Tumour diagnoses were based either on external assessment (suspected), or on histopathology, cytogenetics or previous microsatellite genotyping (confirmed).

## References

Alizon, S., Hurford, A., Mideo, N., and Van Baalen, M. (2009). Virulence evolution and the trade-off hypothesis: history, current state of affairs and the future. J Evol Biol 22, 245–259.

Covarrubias-Pazaran, G., Diaz-Garcia, L., Schlautman, B., Salazar, W., and Zalapa, J. (2016). Fragman: an R package for fragment analysis. BMC Genetics 17, 62.

Deakin, J.E., Bender, H.S., Pearse, A.M., Rens, W., O’Brien, P.C., Ferguson-Smith, M.A., Cheng, Y., Morris, K., Taylor, R., Stuart, A., et al. (2012). Genomic restructuring in the Tasmanian devil facial tumour: chromosome painting and gene mapping provide clues to evolution of a transmissible tumour. PLoS Genet 8, e1002483.

Hawkins, C.E., Baars, C., Hesterman, H., Hocking, G.J., Jones, M.E., Lazenby, B., Mann, D., Mooney, N., Pemberton, D., Pyecroft, S., et al. (2006). Emerging disease and population decline of an island endemic, the Tasmanian devil Sarcophilus harrisii. Biological Conservation 131, 307–324.

Koo, C., Malapi-Wight, M., Kim, H.S., Cifci, O.S., Vaughn-Diaz, V.L., Ma, B., Kim, S., AbdelRaziq, H., Ong, K., Jo, Y.K., et al. (2013). Development of a real-time microchip PCR system for portable plant disease diagnosis. PLoS One 8, e82704.

Lazenby, B.T., Tobler, M.W., Brown, W.E., Hawkins, C.E., Hocking, G.J., Hume, F., Huxtable, S., Iles, P., Jones, M.E., Lawrence, C., et al. (2018). Density trends and demographic signals uncover the long‐term impact of transmissible cancer in Tasmanian devils. Journal of Applied Ecology 0, 1–12.

Loh, R., Bergfeld, J., Hayes, D., O’Hara, A., Pyecroft, S., Raidal, S., and Sharpe, R. (2006). The pathology of devil facial tumor disease (DFTD) in Tasmanian Devils (Sarcophilus harrisii). Veterinary pathology 43, 890–895.

McCallum, H. (2008). Tasmanian devil facial tumour disease: lessons for conservation biology. Trends in ecology & evolution 23, 631–637.

Metzger, M.J., Reinisch, C., Sherry, J., and Goff, S.P. (2015). Horizontal Transmission of Clonal Cancer Cells Causes Leukemia in Soft-Shell Clams. Cell 161, 255–263.

Metzger, M.J., Villalba, A., Carballal, M.J., Iglesias, D., Sherry, J., Reinisch, C., Muttray, A.F., Baldwin, S.A., and Goff, S.P. (2016). Widespread transmission of independent cancer lineages within multiple bivalve species. Nature 534, 705–709.

Murchison, E.P. (2008). Clonally transmissible cancers in dogs and Tasmanian devils. Oncogene 27 Suppl 2, S19–30.

Murchison, E.P., Schulz-Trieglaff, O.B., Ning, Z., Alexandrov, L.B., Bauer, M.J., Fu, B., Hims, M., Ding, Z., Ivakhno, S., Stewart, C., et al. (2012). Genome sequencing and analysis of the Tasmanian devil and its transmissible cancer. Cell 148, 780–791.

Murchison, E.P., Tovar, C., Hsu, A., Bender, H.S., Kheradpour, P., Rebbeck, C.A., Obendorf, D., Conlan, C., Bahlo, M., Blizzard, C.A., et al. (2010). The Tasmanian devil transcriptome reveals Schwann cell origins of a clonally transmissible cancer. Science 327, 84–87.

Pearse, A.M., and Swift, K. (2006). Allograft theory: transmission of devil facial-tumour disease. Nature 439, 549.

Pye, R.J., Pemberton, D., Tovar, C., Tubio, J.M., Dun, K.A., Fox, S., Darby, J., Hayes, D., Knowles, G.W., Kreiss, A., et al. (2016). A second transmissible cancer in Tasmanian devils. Proc Natl Acad Sci U S A 113, 374–379.

Siddle, H.V., Kreiss, A., Tovar, C., Yuen, C.K., Cheng, Y., Belov, K., Swift, K., Pearse, A.M., Hamede, R., Jones, M.E., et al. (2013). Reversible epigenetic down-regulation of MHC molecules by devil facial tumour disease illustrates immune escape by a contagious cancer. Proc Natl Acad Sci U S A 110, 5103–5108.

Stammnitz, M.R., Coorens, T.H., Gori, K.C., Hayes, D., Beiyuan, F., Wang, J., Martin-Herranz, D.E., Alexandrov, L.B., Baez-Ortega, A., Barthorpe, S., et al. (2018). The origins and vulnerabilities of two transmissible cancers in Tasmanian devils.Cancer Cell. https://doi.org/10.1016/j.ccell.2018.03.013

Taylor, B.J., Howell, A., Martin, K.A., Manage, D.P., Gordy, W., Campbell, S.D., Lam, S., Jin, A., Polley, S.D., Samuel, R.A., et al. (2014). A lab-on-chip for malaria diagnosis and surveillance. Malar J 13, 179.

Tovar, C., Obendorf, D., Murchison, E.P., Papenfuss, A.T., Kreiss, A., and Woods, G.M. (2011). Tumor-specific diagnostic marker for transmissible facial tumors of Tasmanian devils: immunohistochemistry studies. Vet Pathol 48, 1195–1203.

